# Towards a genome sequence for every animal: where are we now?

**DOI:** 10.1101/2021.08.04.455150

**Authors:** Scott Hotaling, Joanna L. Kelley, Paul B. Frandsen

**Affiliations:** School of Biological Sciences, Washington State University, Pullman, WA, USA; LOEWE Centre for Translational Biodiversity Genomics (LOEWE- TBG), Frankfurt, Germany; Department of Plant and Wildlife Sciences, Brigham Young University, Provo, UT, USA; Data Science Lab, Smithsonian Institution, Washington, DC, USA

**Keywords:** animal genomes, genome biology, Arthropoda, Mammalia, Metazoa, genomic natural history

## Abstract

In less than 25 years, the field of animal genome science has transformed from a discipline seeking its first glimpses into genome sequences across the Tree of Life to a global enterprise with ambitions to sequence genomes for all of Earth’s eukaryotic diversity (1). As the field rapidly moves forward, it is important to take stock of the progress that has been made to best inform the discipline’s future. In this Perspective, we provide a contemporary, quantitative overview of animal genome sequencing. We identified the best available genome assemblies on GenBank, the world’s most extensive genetic database, for 3,278 unique animal species across 24 phyla. We assessed taxonomic representation, assembly quality, and annotation status for major clades. We show that while tremendous taxonomic progress has occurred, stark disparities in genomic representation exist, highlighted by a systemic overrepresentation of vertebrates and underrepresentation of arthropods. In terms of assembly quality, long-read sequencing has dramatically improved contiguity, whereas gene annotations are available for just 34.3% of taxa. Furthermore, we show that animal genome science has diversified in recent years with an ever-expanding pool of researchers participating. However, the field still appears to be dominated by institutions in the Global North, which have been listed as the submitting institution for 77% of all assemblies. We conclude by offering recommendations for how we can collectively improve genomic resource availability and value while also broadening global representation.

**Significance statement:** The field of animal genome science is rapidly developing, and efforts are underway to sequence genomes for all of Earth’s eukaryotic biodiversity. Here, we provide an overview of animal genome sequencing, with emphases on taxonomic representation, assembly quality, and geographic representation. We show that while a staggering 3,278 unique animal species have had their genomes sequenced, massive disparities exist in terms of the taxonomic groups receiving attention, the quality of the resources being produced, and the institutions driving the field. We highlight areas where improvements can be made, notably by continuing to increase the quality of genome assemblies, including by improving metadata and voucher specimen associations, and actively developing meaningful collaborations between researchers form the Global North and South.

## Introduction

The first animal genome sequence was published 23 years ago (2). The 97 million base pair (Mb) *Caenorhabditis elegans* genome assembly ushered in a new era of animal genome biology where genetic patterns and processes could be investigated at genome scales. As genome assemblies have accumulated for an increasingly diverse set of species, so too has our knowledge of how genomes vary and shape Earth’s biodiversity (e.g., 3, 4). Major shifts in genome availability and quality have been driven by two key events. First, the invention of high-throughput, short-read sequencing provided an economical means to generate millions of reads for any species from which sufficient DNA could be obtained. These ~100-bp short reads could be assembled into useful, albeit fragmented, genome assemblies. Later, the rise of long-read sequencing allowed for similarly economical generation of reads that are commonly orders of magnitude longer than short reads, resulting in vastly more contiguous genome assemblies (5).

We have now entered an era of genomic natural history. Building on ~250 years of natural history efforts to describe and classify the morphological diversity of life on Earth, we are gaining a complementary genomic perspective of Earth’s biodiversity. However, a baseline accounting of our progress towards a complete perspective of Earth’s genomic natural history—where every species has a corresponding, reference-quality genome assembly available—has not been presented. This knowledge gap is particularly important given the momentum towards sequencing all animal genomes which is being driven by a host of sequencing consortia. For instance, the Vertebrates Genome Project seeks to generate high-quality assemblies for all vertebrates (6), the Bird10K project seeks to generate assemblies for all extant birds (7), the i5K Project plans to produce 5,000 arthropod genome assemblies (8), the Earth BioGenome Project aims to sequence all eukaryote genomes (1), and the Darwin Tree of Life project plans to sequence all eukaryotes in Britain and Ireland (https://www.darwintreeoflife.org/).

In this perspective, we curated, quantified, and summarized genomic progress for a major component of Earth’s biodiversity: kingdom Animalia (Metazoa) and its roughly 1.66 million described species (9). We show that as of June 2021, 3,278 unique animals have had their nuclear genome sequenced and the assembly made publicly available on the National Center for Biotechnology Information (NCBI) GenBank database (10). This translates to 0.2% of all animal species, but when viewed through the lens of major clades, massive disparities exist. For instance, 32x more assemblies are available for chordates relative to arthropods (Fig. 1).

**Figure 1.**
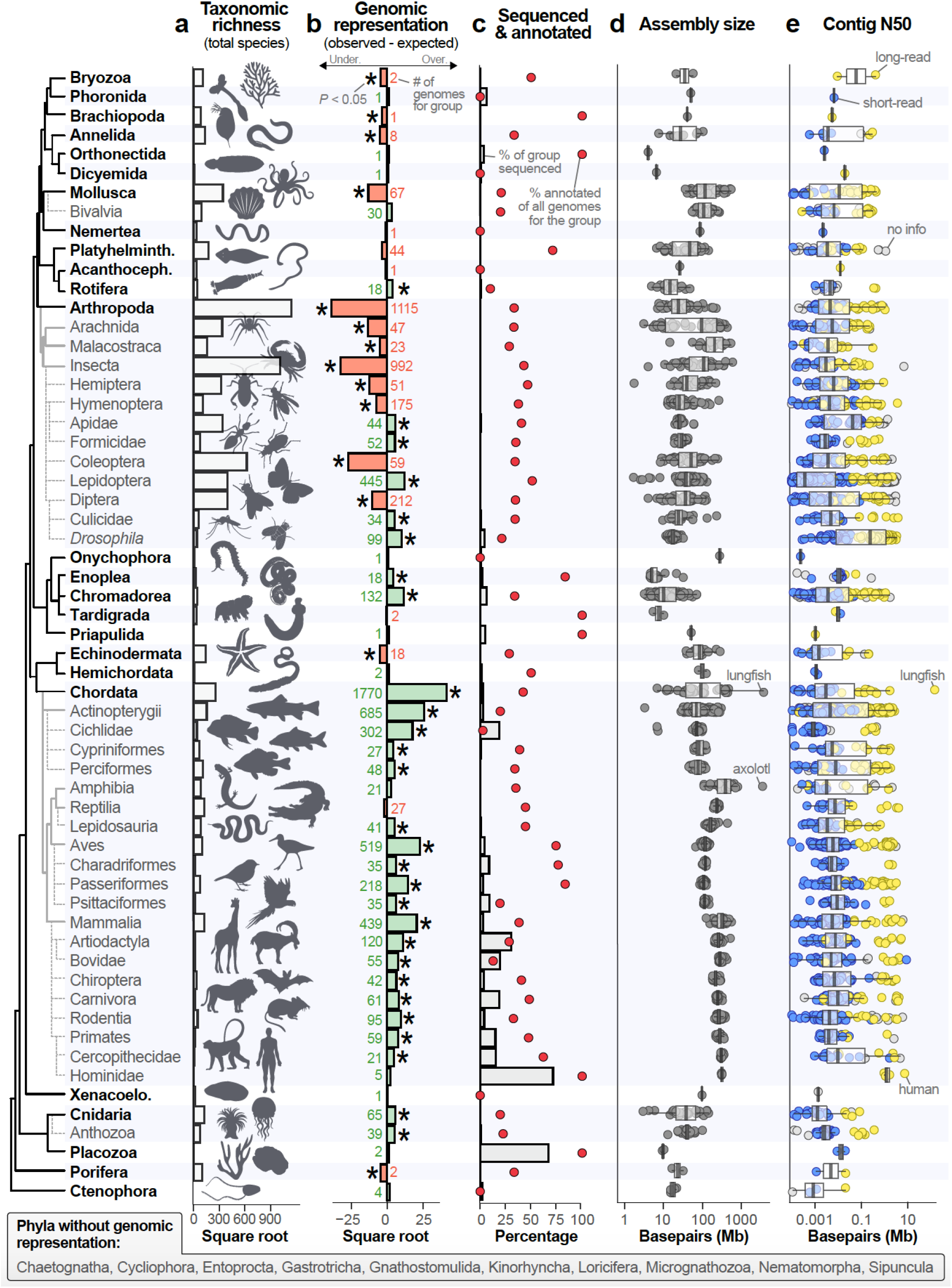
Variation in taxonomic richness and genome availability, quality, and assembly size across kingdom Animalia on GenBank (as of 28 June 2021). Taxonomic groups are clustered by phylogeny following (12). Only groups with 30 or more available assemblies as of January 2021 are shown with the exception of Hominidae (*n* = 5 genomes). In the tree, bold group names represent phyla and naming conventions follow those of the NCBI database. Of 34 recognized animal phyla, 10 do not have a representative genome sequence. (a) The total number of described species for each group following Zhang (9) and the references therein. (b) Genomic representation among animal groups for 3,278 species with available genome assemblies. Bars represent the magnitude of the observed minus the expected number of genomes given the proportion that each group comprises of described animal diversity. Significance was assessed with Fisher’s exact tests and significantly under- or over-represented groups (*P* < 0.05) are denoted with asterisks. Gray numbers indicate the total number of species with available genome assemblies for each group. The number of available assemblies is not mutually exclusive with taxonomy; that is, a carnivore genome assembly would be counted in three categories (Order Carnivora, Class Mammalia, Phylum Chordata). (c) The percentage of described species within a group with an available genome sequence (bars) and the percentage of those assemblies that have corresponding annotations (red circles). For many groups (e.g., arthropods), only a fraction of a percent of all species have an available genome assembly, making their percentage appear near zero. (d) Assembly size for all animal genome assemblies, grouped by taxonomy. (e) Contig N50 by taxonomic group. The sequencing technology used for each assembly is denoted by circle fill color: short-read (blue), long-read (yellow), or not provided (gray). In (d) and (e), each circle represents one genome assembly and a few notable or outlier taxa are indicated with gray text.

## Methods

To construct a database of the best available genome assembly for all animals, we downloaded metadata from GenBank for all kingdom Animalia taxa using the “summary genome” function in v.10.9.0 of the NCBI data sets command line tool on 4 February 2021. Next, we used the TaxonKit (11) “lineage” function to retrieve taxonomic information for each taxid included in the genome metadata. To gather additional data for each assembly (e.g., sequencing technology), we used a custom web scraper script. Both this web scraper script and the scripts used to download and organize the metadata are available on this study’s GitHub repository (https://github.com/pbfrandsen/metazoa_assemblies). We later supplemented this initial data set with a second round of metadata acquisition on 28 June 2021. For the full data set, we hand-refined the NCBI taxonomy classifications to subdivide our data set into three categories: species, subspecies, or hybrids (Table S1). If replicate assemblies for a taxon were present, we defined the “best available” assembly as the one with the highest contig N50 (the mid-point of the contig distribution where 50% of the genome is assembled into contigs of a given length or longer).

We filtered our data in several ways: we removed subspecies (unless they were the only representative for a species) and hybrids, assemblies that were shorter than 15.3 Mb [the smallest confirmed assembly size for a metazoan to date, (13)], or had a contig N50 less than 1 kilobase (Kb). We also culled assemblies that were unusually short (i.e., 1-2.5 Mb) and had information in their descriptions that indicated they were not true nuclear genome assemblies (e.g., “exon capture”). In total, we culled 407 assemblies that did not fit the above criteria. The remaining assemblies were classified as “short-read”, “long-read”, or “not provided” based on whether only short-reads (e.g., Illumina) were used, any long-read sequences (e.g., PacBio) were used, or no information was available. We defined a species as having gene annotations available if any assembly for that taxon also had annotations on GenBank. When the best available assembly did not have annotations included or when multiple assemblies had annotations, we retained the annotations for the assembly with the highest contig N50. Finally, we used the submitting institution for each assembly as a surrogate for the institution that led the genome assembly effort. Using these data, we classified assemblies to a country, region (Africa, Asia, Europe, Middle East, North America, Oceania, South America, Southeast Asia), and the Global North (e.g., Australia, Canada, Europe, USA) or Global South (e.g., Africa, Asia including China, Mexico, Middle East, South America).

To test if clades were under- or overrepresented in terms of genome availability relative to their species richness, we compared the observed number of species with assemblies to the expected total for the group. We obtained totals for the number of described species overall and for each group from previous studies, primarily from Zhang (9) and the references therein. We assessed significance between observed and expected representation with Fisher’s exact tests (alpha = 0.05). We tested for differences in distributions of contig N50 or assembly size between short- and long-read genomes with Welch’s T-tests. For both display (i.e., Figure 1) and analysis, we subdivided the data set into the lowest taxonomic level that still contained 30 or more assemblies as of January 2021 (with the exception of hominids which were given their own category due to their exceptionally high genomic resource quality).

## Results

### Taxonomic representation

Genome assemblies were available for 3,278 species representing 24 phyla, 64 classes, and 258 orders (Fig. 2a, Table S1). The data set was exceptionally enriched for the phylum Chordata (which includes all vertebrates) with 1,770 assemblies for the group (54% of all assemblies) despite chordates comprising just 3.9% of animal species (*P*, Fisher’s < 1e-5; Fig. 1). Conversely, arthropods were underrepresented with 1,115 assemblies (34% of the data set) for a group that comprises 78.5% of animal species (*P*, Fisher’s < 1e-5; Fig. 1). However, not all arthropods were underrepresented; five insect clades were overrepresented [Apidae (bees), Culicidae (mosquitoes), *Drosophila* (fruit flies), Formicidae (ants), and Lepidoptera (butterflies and moths); all *P*, Fisher’s < 1e-3; Fig. 1]. Collectively, of the 59 animal taxonomic groups included in our data set, 14 groups were underrepresented, 17 were represented as expected, and 28 were overrepresented (primarily chordates; Fig. 1). Ten phyla had no publicly available genome sequence (Fig. 1). Over the ~17-year GenBank genome assembly record, animal assemblies have been deposited at a rate of 0.52 species’ assemblies per day. Over the most recent year, however, this rate increased eight-fold to 4.07 assemblies per day. If the most recent rate were maintained, all currently described animals would have a genome assembly available by 3136. To achieve this goal by 2031 instead, an average of 165,614 novel animal genomes would need to be sequenced and assembled each year (~112x faster than the rate for the most recent year).

**Figure 2.**
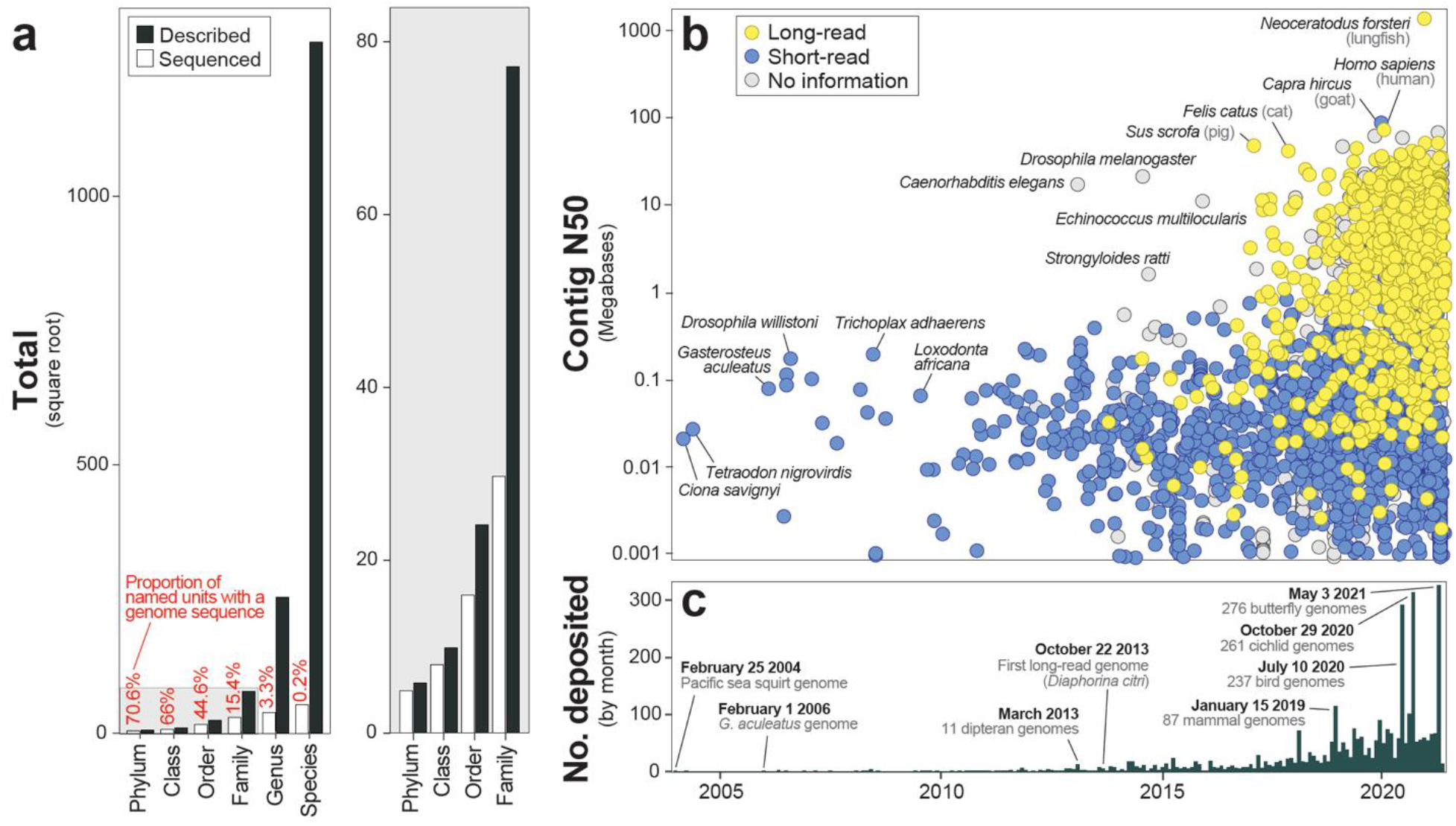
Genome availability for kingdom Animalia versus taxonomic descriptions and over time. (a) The proportion of described taxonomic groups versus the number with sequenced genome assemblies from phyla to species. The gray plot to the right is a zoomed in perspective of the higher taxonomic level categories in the full plot to the left. For genus through phylum, the number of described categories is based on the NCBI taxonomy. For species, the total number described is from Zhang (9). (b) The timeline of genome contiguity versus availability for animals according to the GenBank publication date (x-axis, see panel c). A rise in assembly contiguity has been precipitated by long-read sequencing. Particularly contiguous assemblies for a given time period are labeled. (c) The number of animal genome assemblies deposited on GenBank each month since February 2004. Several notable events are labeled. When specific dates are indicated, those (and the assemblies referred to) are included within that month’s total. For (b) and (c), it is important to note that when a genome assembly is updated to a newer version, its associated date is also updated. Thus, the date associated with many early animal assemblies [e.g., *Caenorhabditis elegans*, (2)] has shifted to be more recent with updates.

### Assembly size, contiguity, and annotations

The average animal genome assembly was 1.02 gigabases (Gb) in length (SD = 1.21 Gb) with a contig N50 of 2.26 Mb (SD = 25.16 Mb; Fig. 1d,e). Two animal genome assemblies were 25 Gb longer than all other assemblies—the axolotl [32.4 Gb (14)] and Australian lungfish [34.6 Gb (15); Fig. 1d]. The smallest genome assembly in the data set, the mite *Aculops lycopersici*, was over 1000x smaller, spanning just 32.5 Mb (16). Still smaller is the 15.3 Mb assembly of the marine parasite, *Intoshia variabili*, which has the smallest animal genome currently known (13). But, since the *I. variabili* assembly was not available on GenBank as of June 2021, it was not included in our data set.

Contiguity varied dramatically across groups. For instance, hominid assemblies (family Hominidae, *n* = 5) were the most contiguous with an average contig N50 of 24.2 Mb. Bird assemblies (class Aves, *n* = 515) were also highly contiguous (mean contig N50 = 1.4 Mb) despite being so numerous (and accumulating for a long period of time). On the other end of the spectrum, jellyfish and related species (phylum Cnidaria) exhibited some of the least contiguous genome assemblies with a mean contig N50 of 0.18 Mb (*n* =65; Fig. 1e). Roughly 34% of animals with genome assemblies had corresponding annotations on GenBank but annotation rates differed substantially among groups (Fig. 1c). For example, the rate of arthropod annotations (22.3%) lags behind that for chordates (41.3%); however much of this disparity appeared to be driven by the low and high annotation rates of butterflies and moths (order Lepidoptera) and birds (class Aves), respectively. Of 445 assemblies, just 6.5% of lepidopteran assemblies on GenBank have corresponding annotations versus 72.8% of birds (*n* = 519 assemblies; Fig. 1c). Notably, since most gene models are based on sequence similarity to known functional genes and not functional data, the true rate of annotation is likely even lower than reported here.

### Geographic representation

Animal genome assemblies have been contributed by researchers at institutions on every continent with permanent inhabitants, including 52 countries. From a regional perspective, institutions in North America (*n =* 1,331), Europe (*n* = 972), and Asia (*n* = 828) collectively accounted for 95.5% of all assemblies (Fig. 3a). And, nearly 70% of all animal genome assemblies have been submitted by researchers in just three countries: USA (*n* = 1,275), China (*n* = 676), and Switzerland (*n =* 317; Fig. 3a). When countries were grouped by their inclusion in the Global North or South, similarly stark patterns emerged. Researchers affiliated with institutions in the Global North contributed roughly 75% of animal genome assemblies (Fig. 3b). From a taxonomic perspective, researchers at North American institutions have contributed the most insect and mammal assemblies, European researchers have contributed the most fish assemblies, and Asian researchers the most bird assemblies (Fig. 3a). The first assembly in GenBank from the Global North was deposited in 2004 and the first assembly from the Global South was deposited in 2011 (Fig. 3c). Since then, the number of assemblies deposited each year has steadily risen with the proportions from the Global North and South staying relatively constant (Fig. 3c).

**Figure 3.**
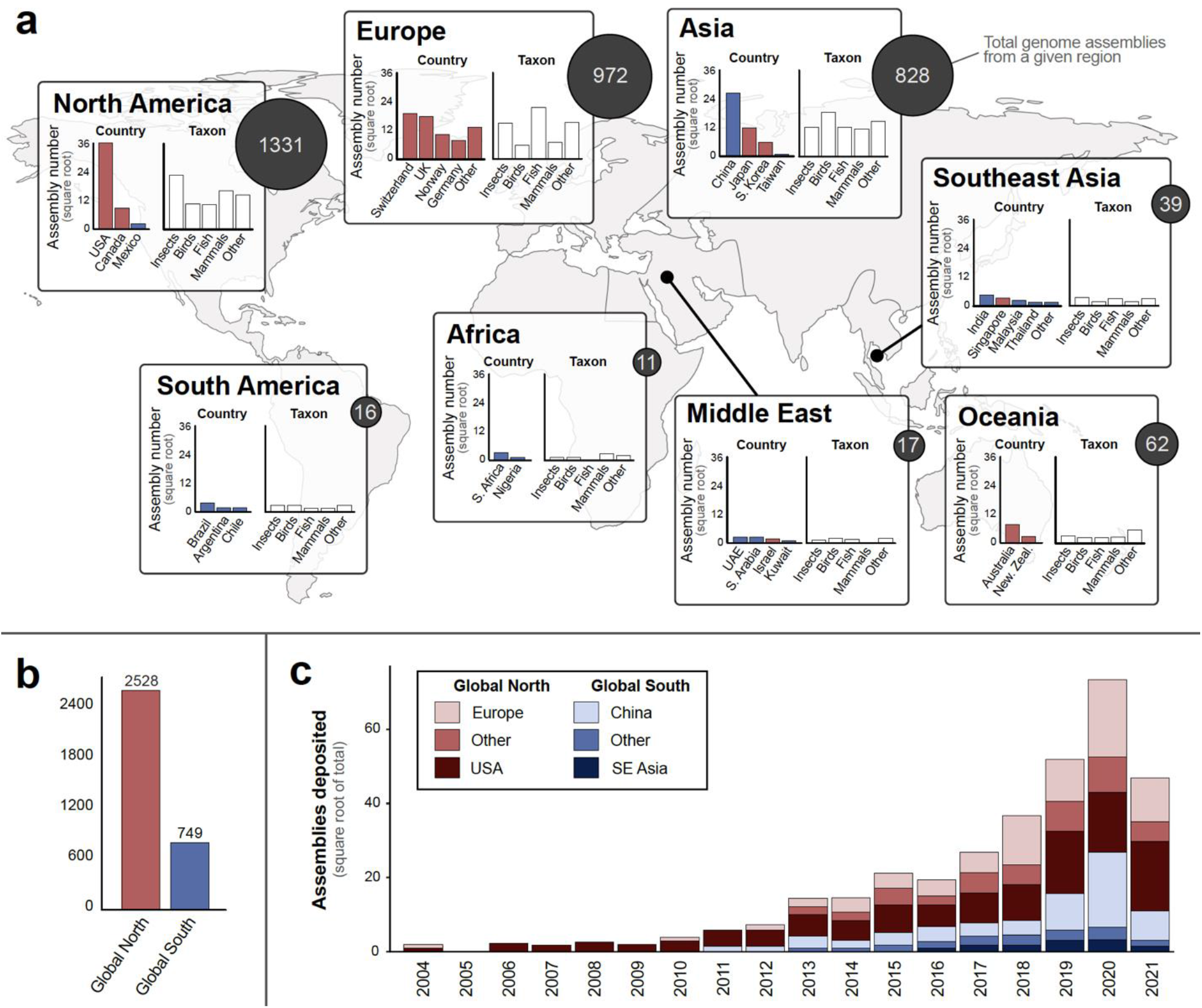
Where animal genome assemblies have been produced around the world according to the submitting institutions on GenBank. (a) For each geographic region, total numbers of genome assemblies are shown by dark circles with white lettering. This total is further broken down by country and taxon. For regions where more than four countries have contributed assemblies (e.g., Europe), an “Other” category represents all other countries. The same applies to all assemblies that are not insects, birds, fish, or mammals in the taxon plots. Countries are color-coded by assignment to the Global North or South. (b) The total number of genome assemblies contributed by countries in the Global North (e.g., USA, Europe, Australia) versus the Global South (e.g., Africa, South America, China, Mexico, Middle East). (c) The rate of genome assembly deposition by major sources in the Global North (Europe, USA) and Global South [China, Southeast (SE) Asia] as well as all other countries collectively in each (Other).

Use of long-reads in genome assemblies and availability of key metadata also differs with geography. For assemblies deposited since 2018, researchers from the Global South have used long-reads slightly more frequently than those from the Global North (25.7% versus 20.2%; Fig. 4a). However, researchers from the Global North were far less likely to report the types of sequence data used (19.9% of assemblies for the Global North versus 1.4% of assemblies for the Global South; Fig. 4a). Much of this difference appears to be driven by genome assemblies deposited by researchers at European institutions (Fig. 4b). This gap in metadata may reflect an issue with data mirroring between the European Nucleotide Archive (ENA) and GenBank. For instance, many new genome assemblies being generated by the United Kingdom, for example, are part of the Wellcome Sanger Institute’s Darwin Tree of Life project, which is generating exceptionally high-quality assemblies using long-read sequencing and depositing them into the ENA (Fig. 5). One region (Oceania) and three countries (Australia, Finland, India) reported long reads being used in more than 50% of deposited assemblies (Fig. 4b,c).

**Figure 4.**
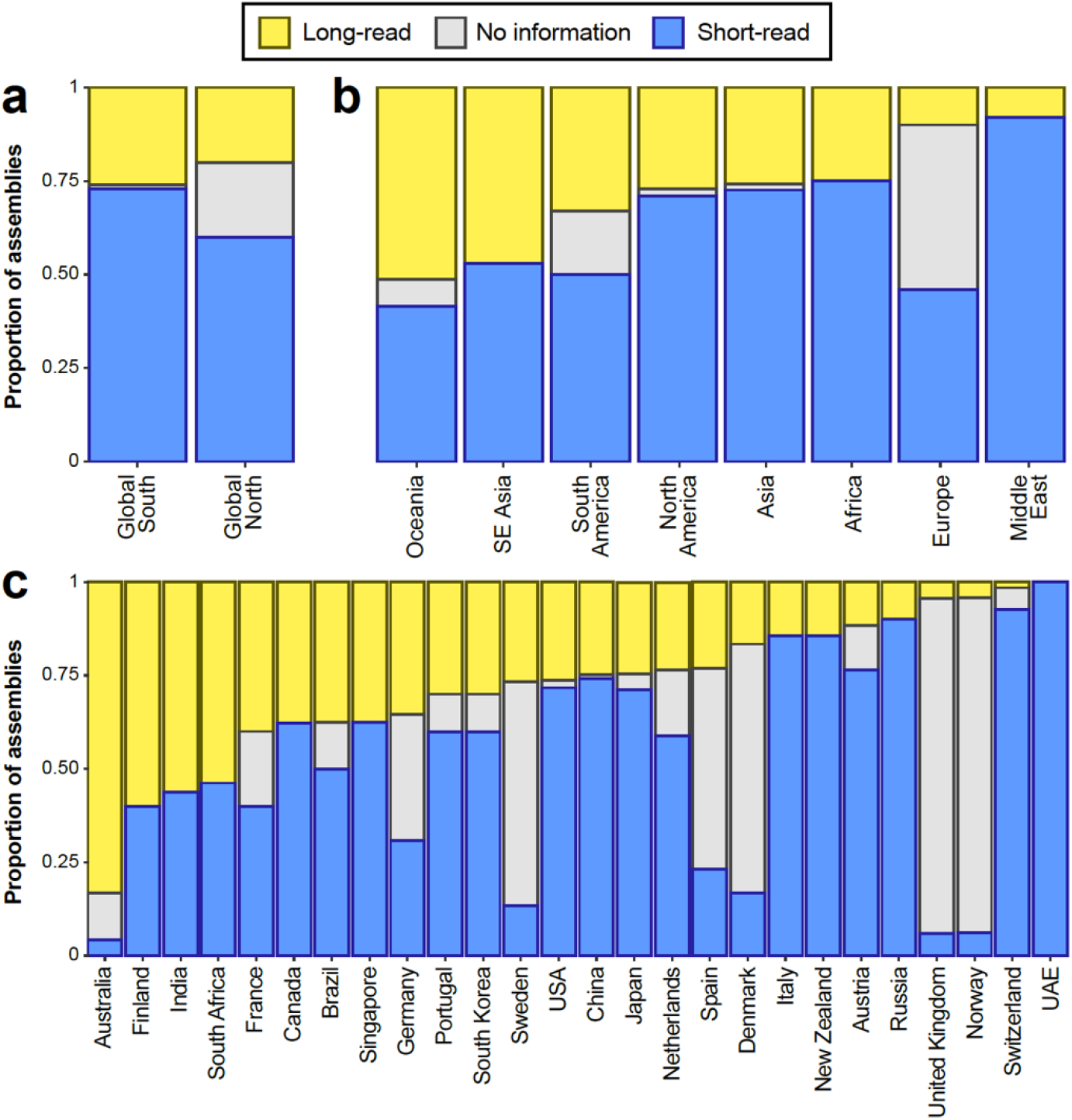
Sequencing technologies used around the world: (a) between the Global North versus Global South, (b) among regions, and (c) among countries. To limit bias due to the limited availability of long-read sequencing technologies before ~2017 (see Fig. 2b), only assemblies deposited on or after 1 January 2018 were included in the analysis and in (c) only countries that deposited five or more assemblies during the focal period (January 2018 – June 2021) are shown.

**Figure 5.**
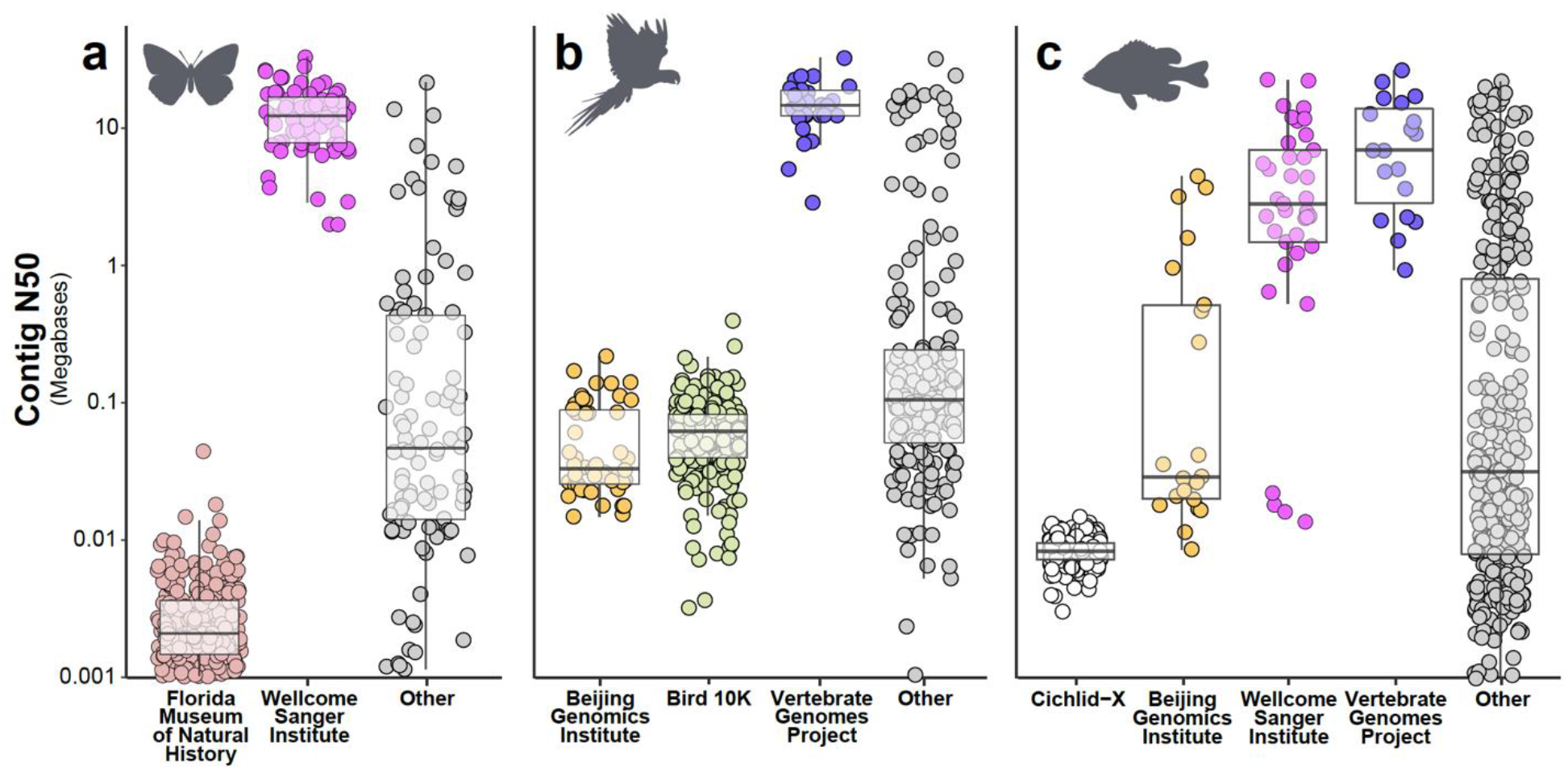
Examples of major contributors of genome assemblies for (a) butterflies (order Lepidoptera), (b) birds (class Aves), and fish (primarily class Actinopterygii). Major contributors were defined as any consortium, organization, or project that has deposited more than 5% of all assemblies for butterflies and birds or 2.5% of all assemblies for fish.

## Discussion

### Taxonomic representation

Animal genome sequencing has dramatically progressed in the last 25 years. In that span, the field has moved from sequencing the first nuclear genome for any animal (2)—a landmark achievement—to targeting the generation of genome assemblies for *all* of Earth’s eukaryotic biodiversity (1). Here, we provided a contemporary perspective on progress towards this goal for the 1.6-million species in the animal kingdom (9). We showed that while tremendous progress has been made, major gaps and biases remain both in terms of taxonomic and geographic representation, at least within the most commonly used database of genomic resources, GenBank. For instance, a major bias exists in favor of vertebrates which are vastly overrepresented relative to their total species diversity (Fig. 1a-c). From the perspectives of biomedicine and human evolution, this bias is reasonable since humans are vertebrates. However, from a basic research perspective, particularly as it relates to genomic natural history and an overarching goal to sequence all animal genomes, there is a need to taxonomically diversify sequencing efforts.

At the highest taxonomic levels, 10 animal phyla still have no genomic representation. To illustrate the scale of this disparity versus other groups and the unique biology that is being overlooked, genome assemblies are available for 685 ray-finned fishes (class Actinopterygii) but none exists for phylum Nematomorpha, a ~2,000-species clade of parasitic worms whose presence can dramatically alter energy budgets of entire stream ecosystems (17). Another phylum without genomic representation—Loricifera—was first described in 1983 (18). This group of small, sediment-dwelling animals includes the only examples of multicellular species that spend their entire life cycles under permanently anoxic conditions (19). Loriciferans accomplish this feat by foregoing the energy-producing mitochondria found in virtually all animals in favor of hydrogenosome-like organelles akin to those found in prokaryotes inhabiting anaerobic habitats (19). Clearly, there is much to discover in terms of genomic diversity and functional biology in undersampled clades.

### Global representation

A few select countries—primarily the USA, several European nations, and China—have led the sequencing of the vast majority of animal genome assemblies (Fig. 3a). Aside from China, all of these countries are within the Global North. This pattern of geographic bias raises two potential issues for representation in animal genome science. First, the researcher population of animal genome sequencing likely does not reflect the global population. Second, sampling biases may exist towards the regions where most of the genome sequencing is occurring. Some of this “bias” is intentional and reflects funding goals for a given region. For instance, the Darwin Tree of Life project (https://www.darwintreeoflife.org/) seeks to sequence the genomes of all ~70,000 eukaryotic species living in Britain and Ireland. Still, however, similar to how sampling biases can yield skewed understanding of the natural world in other disciplines (e.g., 20), so too could bias towards specific ecoregions, habitats, or other classifications skew genomic insight.

Inherently linked with questions of representation in animal genome science is the specter of parachute science (or helicopter research)—the practice where international scientists, typically from wealthy nations, conduct studies in other countries that are often poorer without meaningful communication nor collaborations with local people (21). Parachute science has a long history in ecological research, and signatures of these practices have been observed for genome sciences. For instance, Marks *et al*. (22) found that the majority of plant genome assemblies for species that are native to South America and Africa were sequenced “off-continent” by researchers at European, North American, or Asian institutions. Given the sheer number of animal genome assemblies that have been submitted by a small number of countries and institutions, a similar pattern likely exists for animal genome science. However, to properly assess this issue, parsing authorship to quantify collaboration, at a minimum, would need to occur and this approach would still overlook key aspects of representation that need to be considered (e.g., if a researcher from the Global South is working at an institution in the Global North).

### Assembly quality varies dramatically within and among clades

For the purpose of biological discovery, not all genome assemblies are created equal. As long-read sequencing technologies have matured, so too have the quality of assemblies being generated (5). In the last year alone, the largest ever animal genome assembly was deposited [Australian lungfish, (15)] as well as the most complete human genome to date, a telomere-to-telomere assembly (23). Still, many species on GenBank only have low-quality assemblies available (i.e., contig N50 < 100 Kb with no corresponding gene annotations; Fig. 1). Since fragmentation and/or poor or missing gene annotations reduces the research value of an assembly, genome quality is an important consideration, particularly when the end goal is resource development for a broader community. As of April 2021, the Earth BioGenome Project sought assembly quality of 6.C.Q40 (https://www.earthbiogenome.org/assembly-standards) for reference genomes where “6” refers to a 1e6 contig N50 (i.e., 1 Mb). In our data set, 568 assemblies (17.3%) reach this contiguity standard. And, that number drops to 271 assemblies (8.3%) when contig N50 ≥ 1 Mb and deposited gene annotations are both required. For reference, the “C” above refers to chromosomal scale scaffolding and “Q40” to a less than 1/10,000 error rate. Neither of these metrics were assessed in this study.

Independent research laboratories, institutions, and consortia have contributed genome assemblies on both ends of the quality spectrum (Fig. 5). For example, among butterflies (order Lepidoptera), a bimodal quality distribution is being primarily driven by contributions made in 2021 by two submitting institutions, the Florida Museum of Natural History (e.g., 24) and the Wellcome Sanger Institute (Fig. 5a). When viewing genome assembly contributions holistically across the animal Tree of Life, it is clear that two consortia—the Vertebrate Genomes Project (6) and the Darwin Tree of Life (https://www.darwintreeoflife.org/), part of the Wellcome Sanger Institute—warrant specific recognition for contributing exceptional genomic resources relative to closely related species (Fig. 5a-c).

### Going forward

While animal genome science has dramatically matured in recent years, the field still rests on the cusp of massive change. Thousands of genome assemblies are now available for a wide range of taxa, a resource that can empower unprecedented scales of genomic comparison. Simultaneously, multiple consortia are building momentum towards their goals and generating some of the highest quality genome assemblies ever produced. The field is also diversifying with researchers around the world, particularly from the Global South, leading a rising number of efforts. These ongoing advances will yield higher quality, more globally representative genomic data for animals. As we collectively build towards this new genomic future, we offer recommendations to improve assembly quality and accessibility while also continuing to increase representation within the discipline.

The quality of a genome assembly is likely the most important factor dictating its long-term value. Genome assembly “quality,” however, is difficult to define. Here, we propose a holistic view on genome assembly quality that generally echoes the guidelines proposed by the Earth BioGenome Project and other consortia. Briefly, assemblies should reach minimum levels of contiguity (e.g., contig N50 > 1 Mb) and accuracy in order to be considered a reference that will likely not need to be updated for most applications. At a minimum, assemblies should also include high-quality gene annotations that perhaps take advantage of standardized pipelines [e.g., NCBI Eukaryotic Genome Annotation Pipeline, (25)] to maximize compatibility across taxa. We recommend the field further improve the quality of genome assembly resources in two ways. First, refining and expanding the coordinated deposition of genome assemblies will improve the usability of the resources and reproducibility of analyses. It will also reduce duplications of effort—i.e., when a group sequences a genome that has already been produced—an issue that is likely to become increasingly common.

To refine and expand coordinated resource deposition, we recommend the continued use of GenBank (10) or one of the other archives that are members of the International Nucleotide Sequence Database Collaboration (INSDC)—the European Nucleotide Archive (ENA) and DNA Database of Japan (DDBJ)—as the central repositories for genome assemblies and their metadata given their tripartite data sharing agreement. Next, we call on genetic archive administrators, consortia, and independent researchers to collectively improve the metadata submitted with each assembly and the mirroring of data across repositories. Too many assemblies lack basic information about the sequence data and methods used (e.g., Fig. 4), and with the difficulty of linking assemblies to published studies (if available), it can be challenging or impossible to find this information. Further, an expansion of the metadata associated with each assembly—ideally to make more of the categories required and expand demographic data—would make efforts to quantify geographic representation, for instance, far more straightforward. Alternatively, the metadata associated with genome assembly accessions could be integrated with existing efforts like the Genomic Observatories Metadatabase (GeOMe; 26). Furthermore, a set of minimum quality characteristics for a genome assembly may need to be defined. A number of exceptionally low-quality “genome assemblies” (e.g., with contig N50 values shorter than 1 Kb) that often cover only a small fraction of the expected total genome sequence length for a given group are present on GenBank. The presence of these assemblies raises the question: where is the inflection point between resource quality and value to other researchers versus diluting the resources of a shared repository?

For our second recommendation, we amplify and expand the message of Buckner *et al*. (27) and Thompson *et al*. (28): genome science needs specimen vouchers. Vouchers serve as a key physical link between taxonomy and molecular insight. Rarely, however, are vouchers referenced in publications of genome assemblies; only 11% of vertebrate assemblies included such a reference as of January 2020 (27). While vouchers represent a physical reference for assessing taxonomic classification or morphological variation, a properly stored voucher could also provide a long-term source of material for future resource improvement. If a physical specimen cannot be deposited, photographs and/or genomic DNA should be deposited in its place (e.g., 29). Tied to the metadata discussion above, additional fields should be added to GenBank genome assembly accessions to directly link the assembly to a specimen, photos, or genomic DNA that has been deposited elsewhere.

Though geographic representation in animal genome science has improved in recent years, the discipline appears far from properly reflecting the global researcher pool. This issue is almost certainly multi-faceted, likely stemming from a lack of infrastructure (e.g., fewer high-throughput sequencing platforms in developing countries), fewer resources for expensive molecular research, and a corresponding lack of training in genome data analysis. To bridge this gap and to empower a more diverse discipline, the nations and institutions that are devoting large amounts of resources to animal genome sequencing (e.g., China, United Kingdom, USA), and the researchers within those countries, should develop meaningful collaborations with researchers within the countries where their focal species reside (30). These meaningful collaborations—where all parties are valued for their expertise and involved in decision-making—improve the science through transfer of local knowledge, provide a means for local researchers to expand their skillset and network while raising their scholarly profile, and most importantly, can effectively end the practice of parachute science (30). Within-continent (or country) initiatives also have transformative potential for people and genome research. For instance, the African-led effort to sequence three million African genomes over the next 10 years (the “3MAG” project) will yield massive investment in African genomics, an incredible resource for understanding the full scope of human genetic diversity, and a new generation of African genome scientists (31). While focused on human genetics, the infrastructure and expertise that arise from the 3MAG project will no doubt translate to other taxa in the coming years.

A practical justification also exists for increasing representation in genome science, particularly as we seek to generate genome assemblies for every animal on Earth. The Global South is home to the bulk of the world’s biodiversity (32) and as such, researchers in these regions have greater access to key habitats and specimens. Thus, it behooves everyone, including researchers in the Global North, to deepen collaborations with peers in the Global South while also helping to build indigenous capacity for collection, storage, and sequencing of new specimens.

## Conclusion

Animal genome science continues to grow and expand at an exceptional rate. The coming years will surely see thousands, and perhaps tens of thousands, of new genome assemblies from across the Tree of Life, massive technological and analytical improvements, and some of the largest scale and most in-depth studies of animal genome biology conducted to date. However, if we are to realize the ambitious goals of efforts like the Earth BioGenome Project—a self-described biological “moonshot”—the rate and mean quality of animal genome assembly production will have to increase by roughly two orders of magnitude. Regardless of rates and timelines, however, perhaps the most important goal for the future of animal genome science is that we empower a more diverse, representative researcher community in parallel with the generation of new resources.

## Acknowledgements

S.H. and J.L.K. were supported by NSF award OPP-1906015. We would like to thank Guangfeng Song, Eric Cox, and Anne Ketter from the “datasets” development team at the NCBI for their responsiveness and receptiveness to improving this valuable tool for data science.

## References

1. H. A. Lewin et al., Earth BioGenome Project: Sequencing life for the future of life. Proceedings of the National Academy of Sciences 115, 4325–4333 (2018).

2. C. elegans Sequencing Consortium, Genome sequence of the nematode C. elegans: a platform for investigating biology. Science 282, 2012–2018 (1998).

3. C. Hoencamp et al., 3D genomics across the tree of life reveals condensin II as a determinant of architecture type. Science 372, 984–989 (2021).

4. G. W. Thomas et al., Gene content evolution in the arthropods. Genome Biology 21, 1–14 (2020).

5. S. Hotaling et al., Long-reads are revolutionizing 20 years of insect genome sequencing. Genome Biology and Evolution (2021).

6. A. Rhie et al., Towards complete and error-free genome assemblies of all vertebrate species. Nature 592, 737–746 (2021).

7. G. Zhang, Bird sequencing project takes off. Nature 522, 34–34 (2015).

8. i5K Consortium, The i5K Initiative: advancing arthropod genomics for knowledge, human health, agriculture, and the environment. Journal of Heredity 104, 595–600 (2013).

9. Z.-Q. Zhang, Animal biodiversity: An outline of higher-level classification and survey of taxonomic richness (Magnolia press, 2011).

10. E. W. Sayers et al., GenBank. Nucleic Acids Research 48, D84–D86 (2021).

11. W. Shen, J. Xiong, TaxonKit: a cross-platform and efficient NCBI taxonomy toolkit. Biorxiv, 513523 (2019).

12. C. E. Laumer et al., Revisiting metazoan phylogeny with genomic sampling of all phyla. Proceedings of the royal society B 286, 20190831 (2019).

13. G. S. Slyusarev, V. V. Starunov, A. S. Bondarenko, N. A. Zorina, N. I. Bondarenko, Extreme genome and nervous system streamlining in the invertebrate parasite Intoshia variabili. Current Biology 30, 1292–1298. e1293 (2020).

14. S. Nowoshilow et al., The axolotl genome and the evolution of key tissue formation regulators. Nature 554, 50–55 (2018).

15. A. Meyer et al., Giant lungfish genome elucidates the conquest of land by vertebrates. Nature 590, 284–289 (2021).

16. R. Greenhalgh et al., Genome streamlining in a minute herbivore that manipulates its host plant. Elife 9, e56689 (2020).

17. T. Sato et al., Nematomorph parasites drive energy flow through a riparian ecosystem. Ecology 92, 201–207 (2011).

18. R. M. Kristensen, An introduction to loricifera, cycliophora, and micrognathozoa. Integrative and Comparative Biology 42, 641–651 (2002).

19. R. Danovaro et al., The first metazoa living in permanently anoxic conditions. BMC biology 8, 1–10 (2010).

20. A. C. Hughes et al., Sampling biases shape our view of the natural world. Ecography (2021).

21. P. V. Stefanoudis et al., Turning the tide of parachute science. Current Biology 31, R184–R185 (2021).

22. R. A. Marks, S. Hotaling, P. B. Frandsen, R. VanBuren, Lessons from 20 years of plant genome sequencing: an unprecedented resource in need of more diverse representation. bioRxiv (2021).

23. S. Nurk et al., The complete sequence of a human genome. bioRxiv (2021).

24. E. A. Ellis, C. G. Storer, A. Y. Kawahara, De novo genome assemblies of butterflies. GigaScience 10, giab041 (2021).

25. F. Thibaud-Nissen et al., The NCBI Eukaryotic Genome Annotation Pipeline. Journal of Animal Science 94, 184–184 (2016).

26. J. Deck et al., The Genomic Observatories Metadatabase (GeOMe): A new repository for field and sampling event metadata associated with genetic samples. PLoS biology 15, e2002925 (2017).

27. J. C. Buckner, R. C. Sanders, B. C. Faircloth, P. Chakrabarty, Science Forum: The critical importance of vouchers in genomics. Elife 10, e68264 (2021).

28. C. W. Thompson et al., Preserve a voucher specimen! The critical need for integrating natural history collections in infectious disease studies. Mbio 12, e02698–02620 (2021).

29. S. Hotaling et al., Species discovery and validation in a cryptic radiation of endangered primates: coalescent-based species delimitation in Madagascar’s mouse lemurs. Molecular ecology 25, 2029–2045 (2016).

30. F. Adame, Meaningful collaborations can end’helicopter research’. Nature (2021).

31. A. Wonkam (2021) Sequence three million genomes across Africa. (Nature Publishing Group).

32. R. Dirzo, P. H. Raven, Global state of biodiversity and loss. Annual review of Environment and Resources 28, 137–167 (2003).

